# Survivin modulates stiffness-induced vascular smooth muscle cell motility

**DOI:** 10.1101/2024.12.11.628062

**Authors:** Thomas Mousso, Kalina Rice, Bat-Ider Tumenbayar, Khanh Pham, Yuna Heo, Su Chin Heo, Kwonmoo Lee, Andrew T Lombardo, Yongho Bae

**Affiliations:** Department of Pathology and Anatomical Sciences, Jacobs School of Medicine and Biomedical Sciences, University at Buffalo, NY 14203, USA; Department of Pharmacology and Toxicology, Jacobs School of Medicine and Biomedical Sciences, University at Buffalo, Buffalo, NY 14203, USA; Department of Biomedical Engineering, School of Engineering and Applied Sciences, University at Buffalo, Buffalo, NY 14260, USA; Department of Orthopedic Surgery, Perelman School of Medicine, University of Pennsylvania, Philadelphia, PA 19104, USA; Vascular Biology Program, Boston Children’s Hospital, Boston, MA 02115, USA; Department of Biochemistry, Jacobs School of Medicine and Biomedical Sciences, University at Buffalo, Buffalo, NY 14203, USA

## Abstract

Arterial stiffness is a key contributor to cardiovascular diseases, including atherosclerosis, restenosis, and coronary artery disease, it has been characterized to be associated with the aberrant migration of vascular smooth muscle cells (VSMCs). However, the underlying molecular mechanisms driving VSMC migration in stiff environments remain incompletely understood. We recently demonstrated that survivin, a member of the inhibitor of apoptosis protein family, is highly expressed in both mouse and human VSMCs cultured on stiff polyacrylamide hydrogels, where it modulates stiffness-mediated cell cycle progression and proliferation. However, its role in stiffness-dependent VSMC migration remains unknown. To assess its impact on migration, we performed time-lapse video microscopy on VSMCs seeded on fibronectin-coated soft and stiff polyacrylamide hydrogels, mimicking the physiological stiffness of normal and diseased arteries, with either survivin inhibition or overexpression. We observed that VSMC motility increased under stiff conditions, while pharmacologic or siRNA-mediated inhibition of survivin reduced stiffness-stimulated migration to rates similar to those observed under soft conditions. Further investigation revealed that cells on stiff hydrogels exhibited greater directional movement and robust lamellipodial protrusion compared to those on soft hydrogels. Interestingly, survivin-inhibited cells on stiff hydrogels showed reduced directional persistence and lamellipodial protrusion compared to control cells. We also examined whether survivin overexpression alone is sufficient to induce cell migration on soft hydrogels, and found that survivin overexpression modestly increased cell motility and partially rescued the lack of directional persistence compared to GFP-expressing control VSMCs on soft hydrogels. In conclusion, our findings demonstrate that survivin plays a key role in regulating stiffness-induced VSMC migration, suggesting that targeting survivin and its signaling pathways could offer therapeutic strategies for addressing arterial stiffness in cardiovascular diseases.

## I. INTRODUCTION

Arterial stiffness accelerates the progression of various cardiovascular diseases (CVDs) and pathologies, including atherosclerosis, stroke, hypertension, and neointimal hyperplasia^1-4^. Within the tunica media of the vessel, vascular smooth muscle cells (VSMCs) sense increased matrix stiffness caused by vascular injury or elevated high blood pressure, undergoing a phenotypic transition from a contractile to a synthetic state. This shift is characterized by abnormal proliferation, increased synthesis of extracellular matrix (ECM) proteins, and aberrant migration toward the tunica intima^5, 6^, contributing to neointima formation. While reducing arterial stiffness mitigates atheroscleorsis^7^ and neointimal hyperplasia^2^, the underlying mechanisms responsible for the abnormal cellular behavior observed in stiffened regions remain unclear. Various CVD medications, including lipid-lowering agents, antiplatelet drugs, and antihypertensive drugs^8^, have been shown to reduce atherosclerotic plaque formation; however, no current therapy specifically targets arterial stiffening or the associated abnormal cellular behaviors. In addition, drug-eluting stents, which reduce restenosis when combined with antiproliferative and antimigratory agents^9^, can increase the risk of thrombosis by delaying the reendothelialization process^10-12^. Therefore, a deeper understanding of the molecular mechanisms through which arterial stiffness contributes to the progression of various CVDs through modulation of VSMC behaviors may lead to a novel therapeutic strategy.

Survivin, also known as Baculoviral IAP Repeat Containing 5 (Birc 5), belongs to the inhibitors of apoptosis (IAPs) protein family. Survivin has been predominately studied in the context of cancer, where it regulates apoptosis, proliferation, and migration^13-16^. In cardiovascular biology, limited studies have identified its role in the progression of neointimal hyperplasia, heart failure, atherosclerosis, and pulmonary arterial hypertension^17-22^. Adenoviral delivery of a functionally defective survivin mutant attenuated neointimal hyperplasia in mouse^23^ and rabbit^17^ vascular injury models, implying its direct role in this process. Furthermore, given the critical role of stiffness in CVDs, we recently observed significantly elevated survivin expression on high pathological stiffness matrices compared to soft, physiological stiffness in mouse and human VSMCs using an *in vitro* stiffness-tunable polyacrylamide hydrogel model^24, 25^. We further found that survivin plays an important role in regulating cell cycle progression, proliferation, intracellular stiffness, and ECM synthesis under stiff conditions—key cellular processes that drive the progression of CVDs. Aberrant cell migration is another crucial component in promoting various CVDs and pathologies. Survivin has been shown to positively regulate migration in aortic endothelial cells^26, 27^, mesenchymal stromal cell^27^, and VSMCs^28^. Notably, one study performed siRNA-mediated knockdown of survivin, which reduced VSMC chemotaxis in a PDGF-driven transwell assay. However, its role in stiffness-mediated VSMC migration remains unknown, so this study aimed to characterize survivin as a potential regulator of this process.

## II. RESULTS

### A. Survivin inhibition reduces both collective and single cell migration on rigid cell culture plates

A previous study found that siRNA-mediated survivin knockdown reduced VSMC chemotaxis in a PDGF-driven transwell assay^28^, suggesting survivin’s role in migration, consistent with earlier cancer studies^29-31^. To further test the effects of survivin on both collective and single cell migration of VSMCs, we used sepantronium bromide (YM155) to inhibit survivin, a small molecule survivin inhibitor that has demonstrated selective reduction of survivin expression in various cell types^32, 33^. We first confirmed that treating VSMCs with concentrations ranging from 0.1 μM to 2 μM for 24 hours effectively reduced survivin levels (**Fig. 1A, B**), consistent with previous findings^24, 25^. We then assessed the impact of survivin inhibition on collective cell migration using a wound-healing assay. VSMCs were cultured to form a monolayer on tissue culture plates, scratched with a micropipette tip, and treated with varying doses of YM155 or DMSO (vehicle control). Images were taken at 0, 12, and 24 hours post-treatment (**Fig. 1C**). VSMCs treated with DMSO consistently exhibited 60% and 100% wound closure after 12 and 24 hours, respectively **(Fig. 1D)**. Meanwhile, cells treated with YM155 showed a dose-dependent decrease in wound closure at both 12 and 24 hours compared to DMSO-treated cells.

**Fig. 1.**
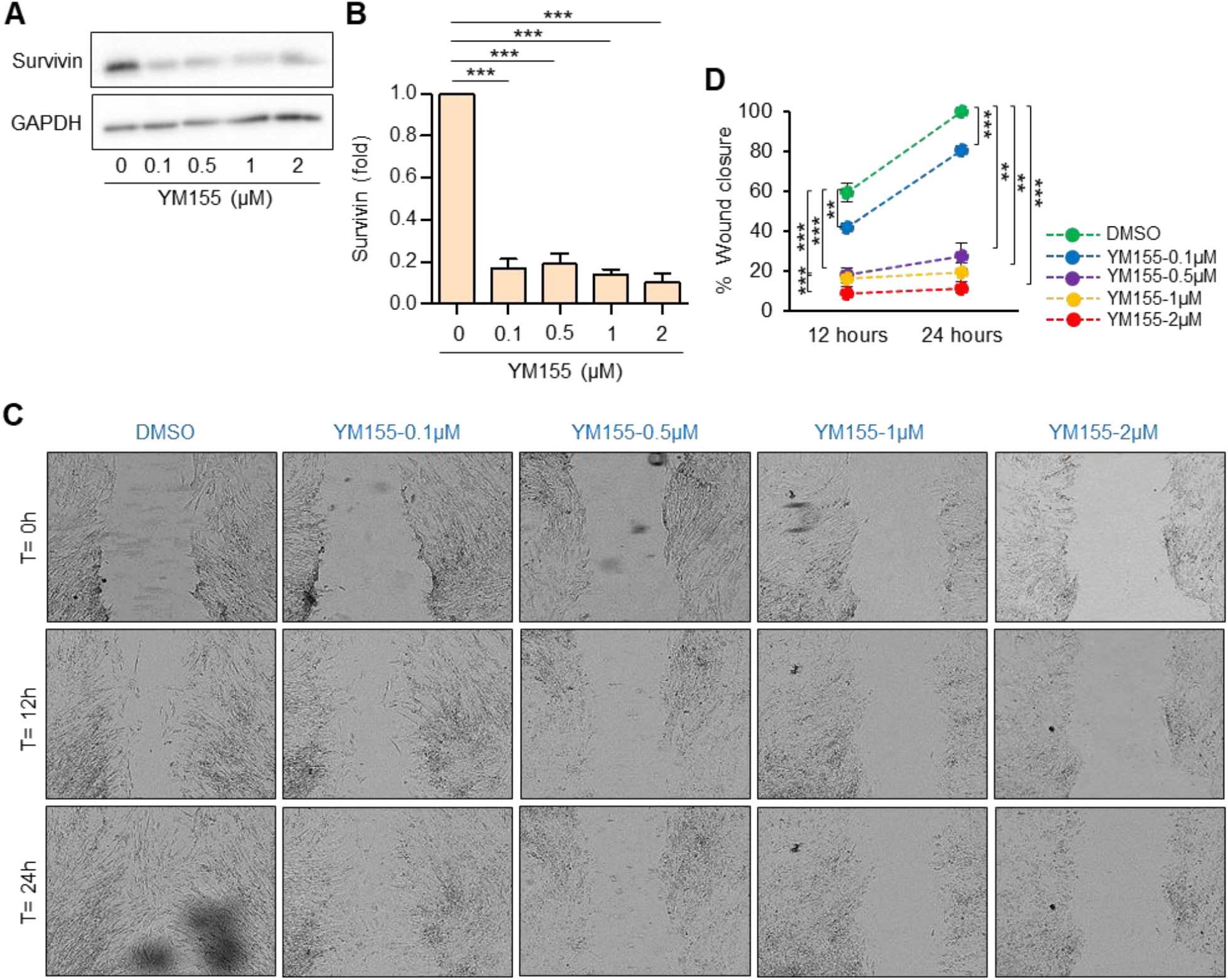
Survivin inhibition reduces collective cell migration. VSMCs plated on cell culture plates were treated with YM155 or DMSO (a vehicle control) at various doses for 24 hours. (**A**) Total cell lysates were collected for immunoblotting. **(B)** The graphs show the expression of survivin in VSMCs treated with YM155, normalized to that in VSMCs treated with DMSO. *n=5*. (**C**) A scratch-wound assay was performed, and images were captured at 0, 12, and 24 hours post-treatment. *n=7-9*. **(D)** Wound closure (%) at 12 and 24 hours was calculated using the formula: [(Initial wound size) - (wound size at 12 or 24 hours)] / (wound size at 12 or 24 hours) *100. *\*\*p<0*.*01, ***p<0*.*001*.

Cell-cell contact can influence the speed of cell migration in the wound-healing assay, potentially limiting the information it provides. To investigate the effect of survivin inhibition on individual VSMCs while minimizing cell-cell contact, we conducted a single-cell analysis of VSMC migration. VSMCs were plated on the tissue culture plates at 10-20% confluency and incubated for 24 hours, followed by treatment with DMSO or varying doses of YM155 for 1 hour. Time-lapse video microscopy was then performed for 3 hours at 3-minute intervals, generating a total of 61 images (frames) for each experimental condition. We first confirmed that YM155 treatment in 4 hours resulted in reduced survivin protein levels in VSMCs **(Fig. 2A, B)**. Using Fiji/Image J software, as described previously^34^, we analyzed the images to track nuclear movement frame-by-frame, allowing us to display the cell trajectory and quantify average velocity. Cell trajectory analysis showed that VSMCs treated with YM155 (0.1, 0.5, 1, and 2 μM) exhibited significantly decreased average single-cell velocities compared to the control (**Fig. 2C**). Interestingly, while observing image sequences of individual cells from different experimental groups treated with YM155 or DMSO, we found that survivin-inhibited cells exhibited multiple short protrusions (loss of directionality), whereas the DMSO-treated control group showed a single prolonged protrusion at the leading edge (**Fig. 2D**). These time-lapse motility observations align with the wound-healing data **(Fig. 1)** and previous transwell migration data^28^, all indicating that survivin is a key modulator of VSMC migration.

**Fig. 2.**
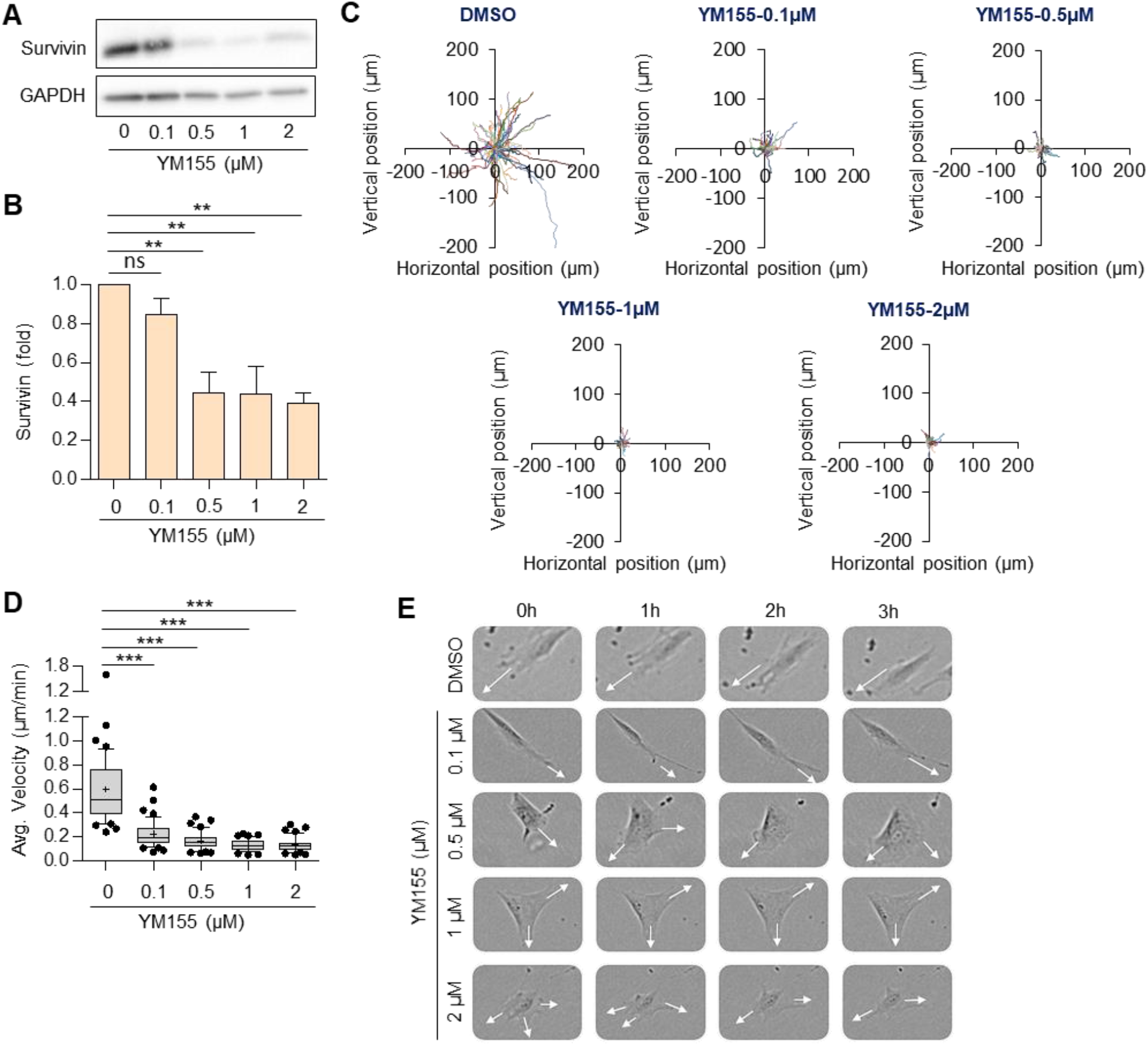
Survivin inhibition decreases single cell migration. (**A**) VMSCs were sparsely plated on cell culture plates and treated with either YM155 or DMSO for 4 hours. Total cell lysates were collected for immunoblotting. **(B)** The graph shows the expression of survivin in VSMCs treated with varying concentrations of YM155, normalized to that of VSMCs treated with DMSO. *n=4*. (**C-E**) VSMCs were plated for time-lapse imaging. Manual tracking of these cells was conducted to obtain single-cell trajectories (**C**) and average cell velocities (**D**). *n=49 (DMSO), n=45 (0*.*1 μM), n=42 (0*.*5 μM), n=44 (1 μM), and n=45 (2 μM)*. (**E**) Sequences of images from a set of representative time-lapse experiments (arrow-direction of protrusion). *\*\*p<0*.*01, ***p<0*.*001*. ns, not significant.

### B. Survivin inhibition reduces stiffness-stimulated cell migration

Although several studies have established substrate stiffness as a critical modulator of cell migration^35-37^, the role of survivin in stiffness-dependent VSMC migration remains unexplored. Here, our results showed that survivin inhibition significantly reduced both collective and single cell migration **(Figs. 1 and 2)**. However, it is important to note that VSMCs were cultured on rigid plastic cell culture dishes, which do not accurately replicate the physiological and pathological stiffness of the *in vivo* environment. To determine whether survivin is required for stiffness-stimulated cell motility, VSMCs were sparsely seeded on fibronectin-coated soft (elastic modulus, 2−8 kPa) and stiff (16−24 kPa) polyacrylamide hydrogels^24, 25^ for 1 hour. The soft hydrogel mimics the physiological stiffness of a healthy mouse femoral artery, while the stiff hydrogel reflects the pathological vessel stiffness seen after vascular injury or atherosclerosis^7, 38^. Cells were then treated with varying doses of YM155 or DMSO, as described above. After 1 hour of treatment, time-lapse video microscopy was performed for 3 hours at 1-minute intervals, yielding 181 images per condition, which were analyzed using Fiji/Image J software. In addition, cell lysates were collected 4 hours after YM155 treatment. We confirmed that survivin expression increases on stiff substrates, as previously shown^24, 25^, and that YM155 reduces survivin levels in VSMCs in a dose-dependent manner, attenuating stiffness-mediated survivin expression (**Fig. 3A, B**). Single cell trajectory analysis revealed that VSMCs on stiff hydrogels exhibited significantly greater migration distance (**Fig. 3C**) and speed (**Fig. 3D**), similar to cells on plastic (**Fig. 2**), compared to those on soft hydrogels. YM155 treatment reduced this increased migration distance and speed to levels similar to those of cells on soft hydrogels. Cells cultured on soft hydrogels displayed a more rounded, less spread morphology with fewer protrusions compared to those on stiff hydrogels (**Fig. 3E**). Similar to cells seeded on plastic (**Fig. 2E**), YM155-treated cells on stiff hydrogels displayed multiple protrusions, while DMSO-treated cells exhibited a single prolonged protrusion at the leading edge (**Fig. 3E**).

**Fig. 3.**
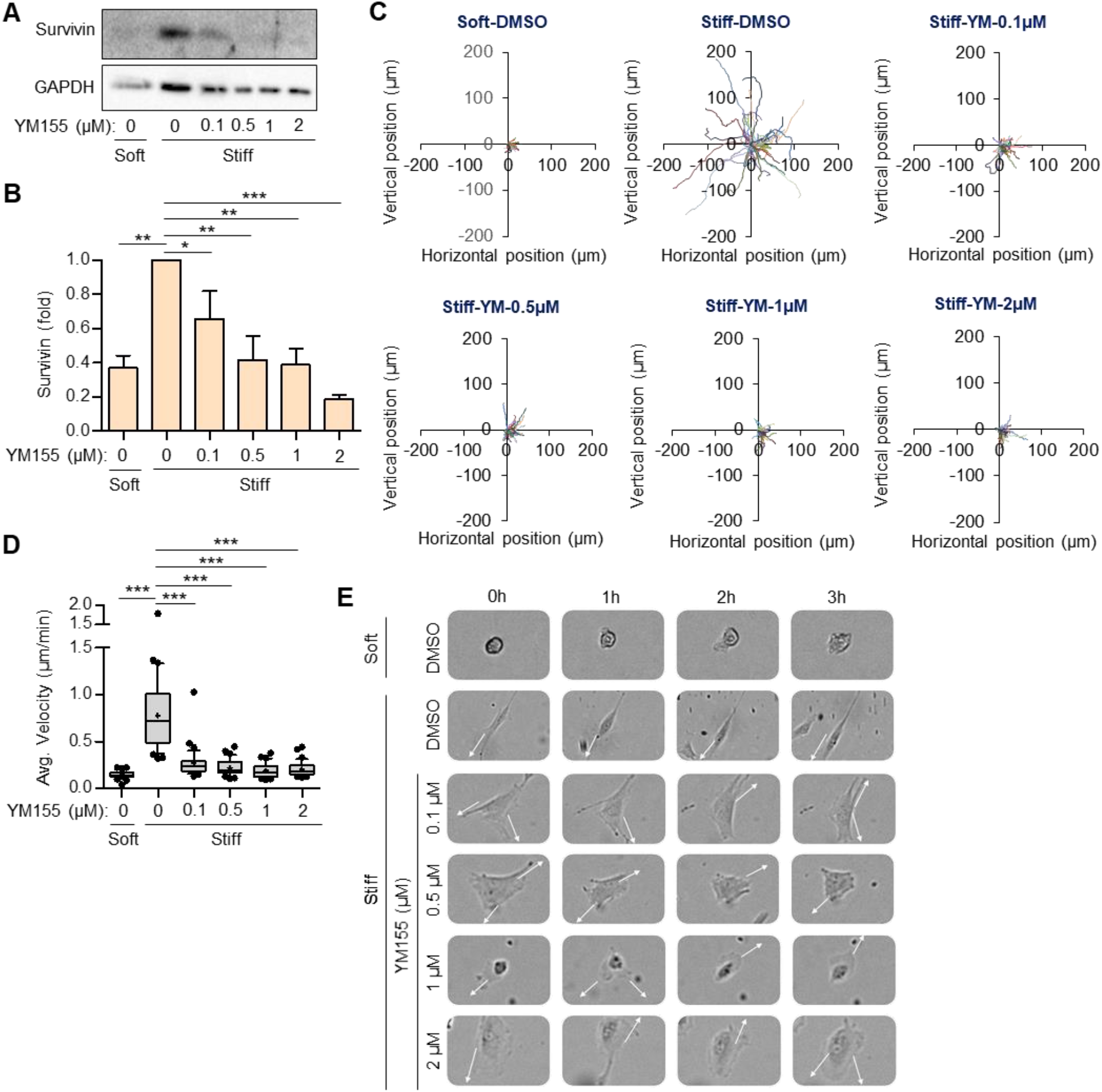
Pharmacological inhibition of survivin reduces stiffness-dependent migration. (**A**) VSMCs were sparsely seeded on fibronectin-coated soft and stiff hydrogels and treated with YM155 or DMSO for 4 hours, followed by collection of total lysates for immunoblotting. **(B)** The graph shows survivin expression in cells treated with varying concentrations of YM155, normalized to the expression in VSMCs on stiff hydrogels treated with DMSO. *n=4*. (**C-E**) VSMCs were plated for time-lapse video microscopy, and manual tracking of these cells was conducted to obtain single-cell trajectories (**C**) and average cell velocities (**D**). *n=35 (Soft-DMSO), n=36 (Stiff-DMSO), n=38 (0*.*1 μM), n=39 (0*.*5 μM), n=35 (1 μM), and n=35 (2 μM)*. (**E**) Sequences of images from a set of representative time-lapse experiments (arrow-direction of protrusion). *\*p<0*.*05, **p<0*.*01, ***p<0*.*001*.

To further confirm whether survivin siRNA-mediated knockdown on cell migration yields similar results to those obtained with YM155, we seeded VSMCs transfected with either survivin siRNA or a non-targeting control siRNA on stiff hydrogels. Survivin siRNA reduced survivin expression in VSMCs compared to cells treated with control siRNA (**Fig. 4A, B**). Similar to YM155, survivin knockdown significantly reduced stiffness-stimulated cell migration (**Fig. 4C, D**) and resulted in multiple protrusions (**Fig. 4E**) compared to cells treated with control siRNA. Collectively, these data from YM155 and siRNA demonstrate that survivin is essential for stiffness-dependent VSMC migration and protrusion.

**Fig. 4.**
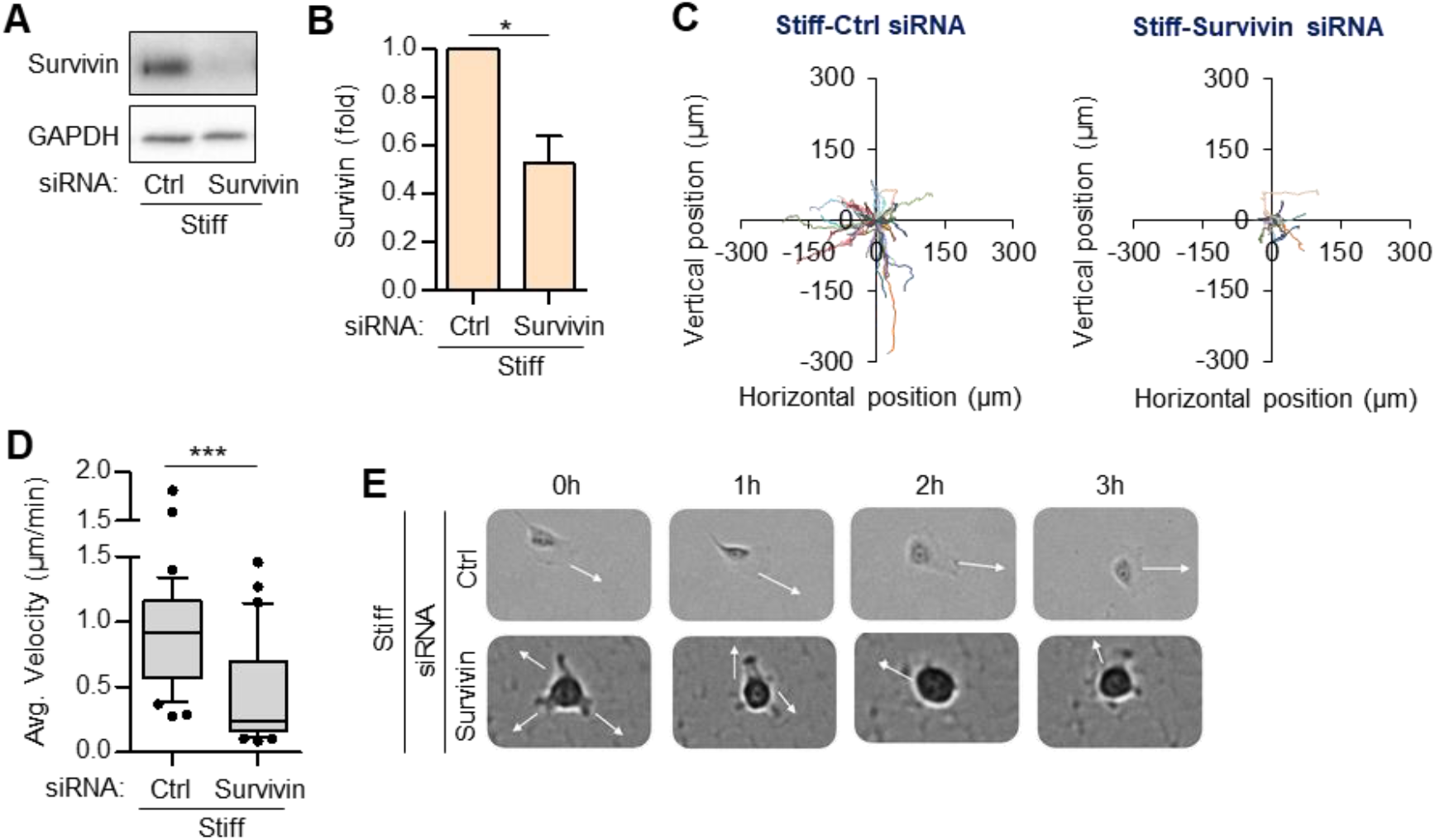
Survivin knockdown decreases stiffness-stimulated cell migration. (**A**) VSMCs transfected with 200 nM survivin siRNA or non-targeting control siRNA were seeded on fibronectin-coated stiff hydrogels for 4 hours. (**B**) Total cell lysates were analyzed by immunoblotting, and survivin protein levels were normalized to GAPDH. *n=3*. (**C-E**) VSMCs were plated for time-lapse video microscopy, and manual tracking of these cells was conducted to obtain single-cell trajectories (**C**) and average cell velocities (**D**). *n=37 (control siRNA) and n=37 (survivin siRNA)*. (**E**) Sequences of images from a set of representative time-lapse experiments (arrow-direction of protrusion). *\*p<0*.*05, ***p<0*.*001*.

### C. Survivin overexpression increases cell migration on soft hydrogels

Survivin protein levels are higher on stiff hydrogels, and cells exhibit greater migratory behavior compared to those on soft hydrogels. To determine whether survivin overexpression alone is sufficient to induce cell migration on soft hydrogels, we infected VSMCs with adenoviruses encoding wild-type survivin or GFP (control) **(Fig. 5A, B)**, and then plated them on soft hydrogels for time-lapse video microscopy. VSMCs infected with adenoviral GFP exhibited a migration rate of 0.169 μm/min. VSMCs treated with adenoviral survivin at 25 and 50 MOI displayed migration rates of 0.249 μm/min and 0.337 μm/min, respectively **(Fig. 5C, D)**. Survivin overexpression increased directional persistence with a prolonged protrusion at the leading edge **(Fig. 5E)**. Collectively, these findings suggest that while survivin partially rescues cell motility and protrusion behaviors under soft conditions, substrate stiffness is still required to fully restore the cell migration observed on stiff conditions.

**Fig. 5.**
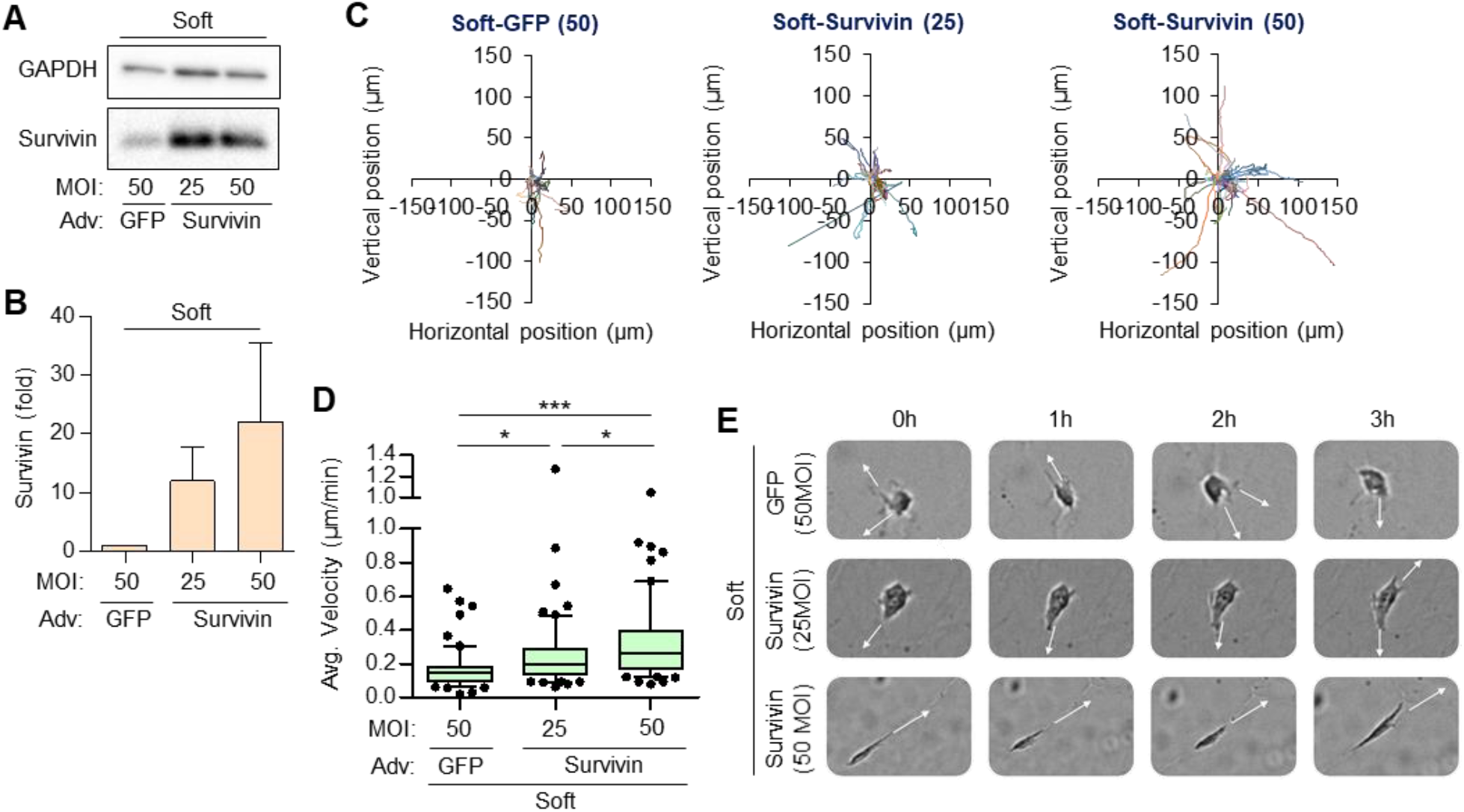
Survivin overexpression partially rescues cell migration on soft hydrogels. (**A**) VSMCs were infected with either adenoviral GFP (50 MOI) or survivin (25 and 50 MOI). (**B**) Total cell lysates were analyzed by immunoblotting, and survivin protein levels were normalized to GAPDH. *n=3*. (**C-E**) VSMCs were plated for time-lapse video microscopy, and manual tracking of these cells was conducted to obtain single-cell trajectories (**C**) and average cell velocities (**D**). *n=59 (GFP 50 MOI), n=60 (survivin 25 MOI), and n=60 (survivin 50 MOI)*. (**E**) Sequences of images from a set of representative time-lapse experiments (arrow-direction of protrusion). *\*p<0*.*05, ***p<0*.*001*. ns, not significant.

## III. DISCUSSION

Survivin has been associated with the progression of neointimal hyperplasia^17, 19^, hypertension^22^, and atherosclerosis^18^, conditions characterized by increased arterial stiffness. Notably, survivin inhibition reduced neointimal hyperplasia in vascular injury models of both mice and rabbits by regulating VSMC migration and proliferation^17, 19^. Recently, we demonstrated that survivin is highly expressed under stiff conditions, where it promotes VSMC proliferation and ECM synthesis^24, 25^. However, its role in stiffness-stimulated VSMC motility remains uncharacterized, and this study is the first to characterize survivin as a key regulator. Our data indicates that stiffness and survivin are critical for cell migration, potentially through increased cell spreading and protrusion―key for extending the leading edge, forming new adhesions to ECM, and enhancing motility^39, 40^. VSMC migration and spreading were increased on stiff hydrogels compared to soft, aligning with observations from Rickel et al.^41^ and our prior study^24^. On soft hydrogels, VSMCs exhibited multiple short protrusions without a defined leading edge, reducing directional migration, whereas on stiff hydrogels, cells displayed a single prolonged protrusion at the leading edge, increasing directional migration. These stiffness-stimulated increases in migration and spreading on stiff hydrogels were significantly attenuated by survivin inhibition, with cells exhibiting multiple short protrusions and reduced cell spreading^24^, similar to those observed on soft hydrogels. Our findings also align with Nabzdyk et al., who showed that survivin modulates VSMC migration, although their study used a transwell assay instead of physiological stiffness conditions^28^. Furthermore, survivin overexpression on soft hydrogels partially rescued migration, with fewer protrusions and increased spreading^24^ compared to wild-type VSMCs on soft hydrogels. This suggests that both stiffness and survivin expression are required for full recovery of VSMC migration observed under stiff conditions.

The mechanism by which survivin regulates stiffness-dependent VSMC migration remains unclear. One potential pathway involves focal adhesion kinase (FAK), which plays a critical role in focal adhesion and actin cytoskeleton organization, both essential for cell motility^42, 43^. Studies have shown that phosphorylation of FAK at Tyr 397 (FAK^pY397^) is necessary for protrusion and migration^44-46^, and is increased under stiff conditions^25, 47^. In prostate cancer cells, survivin inhibition was reported to reduce migration, partially due to decreased FAK^pY397^ levels^48^. This effect was likely driven by reduced recruitment of FAK^pY397^ to focal adhesions complexes leading to diminished cell protrusion. Furthermore, inhibition of FAK^pY397^ was shown to reduce actin stress fiber formation^44^. Stress fibers interact with focal adhesions^49^, establish cell polarity^50^, and play a crucial role in mechanotransduction^51^ by enabling cells to sense matrix stiffness and regulate cell migration^52^. Consistently, a recent study in podocytes demonstrated that survivin inhibition significantly reduced actin stress fiber formation^53^. Based on these findings, we postulated that survivin may regulate stiffness-dependent migration through mechanisms controlling FAK phosphorylation at Tyr397, thereby influencing focal adhesion and actin cytoskeleton organization.

ECM proteins, including collagen-1, lysyl oxidase (Lox), and fibronectin, are also key regulators of cell migration and major contributors to the progression of various CVDs and related pathologies^25, 54^. Differentiated VSMCs synthesize and deposit these ECM proteins, which contribute to arterial stiffening. Our recent study demonstrated a stiffness-dependent increase in collagen-1, fibronectin, and Lox expression and deposition by VSMCs, which was attenuated by survivin inhibition^25^. Other studies have shown that inhibiting newly synthesized collagen in porcine arterial SMCs reduces migration^55^, while fibronectin induces a phenotypic switch that promotes migration^56, 57^. Additionally, Lox inhibition has been shown to reduce atherosclerotic formation and the restenotic process, potentially by decreasing VSMC migration and proliferation^7, 58^. Therefore, another potential mechanism by which survivin may modulate stiffness-dependent VSMC migration is by regulating the expression of ECM proteins.

Our previous study identified survivin as a key mediator of stiffness-dependent VSMC proliferation by regulating the expression of cyclin D1, a major driver of cell cycle progression^24^. Interestingly, a previous study showed that cyclin D1 promotes migration of mouse embryonic fibroblasts by inhibiting Rho-activated kinase signaling^596061^. Additionally, cyclin D1 interacts with and positively regulates filamin A, a cytoskeleton protein involved in cell migration^61^. In pulmonary arterial SMCs, inhibition of filamin A reduces cell motility^62^. Thus, survivin-induced cyclin D1 expression may influence filamin A function, thereby facilitating cell migration.

In summary, understanding the molecular mechanisms driving aberrant cellular behaviors is critical for developing therapeutic strategies for stiffness-associated cardiovascular diseases. Our findings highlight the role of survivin in migration under pathological stiffness conditions, offering insights into the molecular pathways linking ECM stiffness to cardiovascular pathology.

## IV. METHODS

### A. Cell culture

Human vascular smooth muscle cells (VSMCs; catalog number [cat. no.] 354-05a, Cell Applications, Inc.) were cultured in Dulbecco’s modified Eagle’s medium (DMEM; cat. no. 10-014-CV, Corning) supplemented with 10% fetal bovine serum (FBS; cat. no. 2510268RP, GIBCO), 1 mM sodium pyruvate (cat. no. S8636, Sigma), 1% penicillin-streptomycin (cat. no. 30-002-CI, Corning), 50 μg/ml gentamicin (cat. no. 30-005-CR, Corning) and 2% MEM amino acids (cat. no. M5550, Sigma). The cells were maintained at 37°C and 10% CO_2_ and used up to passage 5. To ensure optimal cell growth, the culture medium was replaced every 2−3 days, and the cells were passaged at 80−90% confluency.

### B. Drug Treatment

VSMCs were plated on plastic cell culture plates, glass coverslips, or soft and stiff hydrogels in media containing 10% serum with varying doses (0.1, 0.5, 1, or 2 μM) of YM155 (survivin inhibitor; cat. No. 11490, Cayman Chemical) or dimethyl sulfoxide (DMSO; cat. no. D8418, Sigma) as a vehicle control. VSMCs were incubated with the drug for a predetermined period before being used in assays to assess the impact of YM155 on the cells.

### C. Preparation of polyacrylamide hydrogels

Soft (2−8 kPa) and stiff (18−24 kPa) fibronectin-coated polyacrylamide hydrogels were used to mimic the physiological stiffness of normal (soft) and diseased/injured arteries (stiff), as previously described^25, 38, 63^. Autoclaved glass coverslips were etched with 1.0 M sodium hydroxide for 3 minutes, then treated with 3-(trimethoxysilyl)propyl methacrylate (cat. no. 440159, Sigma-Aldrich) to introduce amine groups for cross-linking with hydrogel. Hydrogel solution was prepared by mixing a solution of a pre-determined ratio^64^ of 40% acrylamide (cat. no. 1610148, Bio-Rad) and 1% bis-acrylamide (cat. no. 1610142, Bio-Rad) with sterile water, 10% ammonium persulfate (cat. no. A3678, Sigma-Aldrich), Tetramethylethylenediamine (TEMED; Cat. No. J63734. AC, Thermo Fisher Scientific), and a solution of N-hydroxysuccinimide (NHS; cat. no. A8060, Sigma-Aldrich)-fibronectin (cat. No. 341631, Calbiochem). The NHS−fibronectin solution was prepared by combining fibronectin (100 μl at 1 mg/ml dissolved in 1.9 ml Tris base solution at pH 8.4) with 222 μl of 1 mg/ml NHS (dissolved in 1 ml of DMSO) and incubated at 37 °C for 1 hour. The solution was then dispensed on the etched/methacrylate-treated glass coverslip, and a siliconized glass coverslip (prepared using 20% Surfasil [cat. no. TS42801, Thermo Scientific] in 80% chloroform [cat. No. J67241.AP, Thermo Scientific]) was placed on top of the dispensed solution. The polymerized hydrogels were extensively washed with Dulbecco’s phosphate-buffered saline (DPBS) to remove unpolymerized polyacrylamide and were blocked with 1 mg/ml heat-inactivated, fatty-acid-free bovine serum albumin (BSA) for 1 hour.

### D. Wound-healing assay

VSMCs were seeded on a cell culture plate, and once the cells reached confluence, a scratch was made down the middle of the monolayer using a p200 micropipette tip. The cells were rinsed once with warm DPBS and incubated in DMEM containing YM155 or DMSO. Images were obtained with a Cytation 1 Imaging Multimode Reader (Agilent Technologies, Inc). To analyze the results of these assays, images at t = 0, t = 12, and t = 24 hours were used. The visible wound in the cell monolayer was manually annotated, and the wound area in each image was measured. The percent of wound closure after 24 hours was subsequently calculated.

### E. Time-lapse single-cell motility assay and image analysis

VSMCs were seeded at 20-30% confluency on cell culture plates or polyacrylamide hydrogels to minimize cell-cell contact and incubated overnight. Following incubation, cells were treated with YM155 or DMSO and immediately placed in a Cytation 1 Imaging Multimode Reader for time-lapse imaging at 37°C and 10% CO_2_. Images were captured every 3 minutes for 3 hours to track cell migration. The time-lapse images were analyzed using Fiji/ImageJ to determine cell migration distance and speed, as previously described^34^. Cell trajectory was constructed by manually marking frame-by-frame the centroid positions (x, y) of cell nuclei.

### F. siRNA transfection and adenovirus infection

VSMCs were transfected with 100 nM survivin (cat. no. AM16704, Ambion) or control siRNA (ID no. 121294, Ambion) using Lipofectamine 3000 reagent (cat. no. L3000-015, Invitrogen) in Opti-MEM reduced serum media (cat. no. 31985-070, Gibco) as previously described^65^. Five hours after siRNA transfection, cells were serum starved in DMEM containing 1 mg/ml BSA for 48 hours, then plated on soft and stiff hydrogels, with experiments performed within 24 hours. Survivin siRNA sequence is 5′-CCACUUCCAGGGUUUAUUCtt-3′. For adenovirus infection, VSMCs were infected for 24 hours with adenoviruses encoding wild-type survivin (multiplicity of infection [MOI], 25 and 50; cat. no. 1611, Vector Biolabs) or GFP (MOI 50; cat. no. 1060, Vector Biolabs), with GFP serving as the experimental control. Following incubation, cells were plated on soft and stiff hydrogels for experiments.

### G. Protein extraction and immunoblotting

Total cell lysates were collected from VSMCs cultured on soft and stiff hydrogels as described previously^66^. Cells were first lysed by placing hydrogels face-down on warm 5X sample buffer (250 mM Tris [pH 6.8], 10% SDS, 50% glycerol, 0.2% bromophenol blue, and 10 mM 2-mercaptoethanol) for 2 minutes at room temperature. For immunoblotting, equal amounts of extracted protein were fractionated on 12% SDS-polyacrylamide gels, and the fractionated proteins were transferred electrophoretically to a polyvinylidene fluoride (PVDF; cat. no. 10026933, Bio-Rad) membrane using the Trans-Blot Turbo Transfer System (Bio-Rad). The PVDF membrane was blocked in 5% milk in TBST (Tris-buffered saline with 0.1% Tween 20 detergent) for 1 hour at room temperature, then incubated with primary antibodies against survivin (1:250; cat. No. 71G4B7, Cell Signaling Technology) and GAPDH (1:10000; cat. No. 60004-1-Ig, Proteintech) diluted in 5% milk in TBST for 2 hours at room temperature, overnight at 4°C, and an additional 2 hours at room temperature followed by a 30-minute wash with TBST. The membranes were then probed with the secondary antibody, HRP-conjugated Goat Anti-Rabbit IgG (H+L) (1:1000; cat. no. SA00001-2, Proteintech), diluted in 5% milk in TBST for 1 hour at room temperature, then washed three times with TBST. Antibody signals were detected using Clarity (cat. no. 170561, Bio-Rad) or Clarity Max (ca. no. 1705062, Bio-Rad) Western ECL substrates on ChemiDoc XRS+ imaging system and band intensity was analyzed using ImageJ.

### H. Statistical Analysis

Data are presented as the mean ± standard error of the mean (SEM). As appropriate, the statistical analysis was performed using a Student’s t-test or a one-way ANOVA. Results with P-values less than 0.05 (*), 0.01 (**), or 0.001 (***) were considered to be statistically significant.

## ACKNOWLEDGMENTS

This work was supported by NIH/NHLBI grant (R01HL163168) to Y.B.

## AUTHOR DECLARATIONS

### Ethical Approval

Ethics approval is not required.

## Author Contributions

**Thomas Mousso**: Conceptualization (lead); Investigation (lead); Formal analysis (supporting); Writing – original draft (equal). **Kalina Rice**: Conceptualization (lead); Investigation (lead); Formal analysis (lead); Writing – original draft (equal). **Bat-Ider Tumenbayar**: Formal analysis (supporting); Writing – review and editing (supporting). **Khanh Pham**: Investigation (supporting); Writing – review and editing (supporting). **Yuna Heo**: Formal analysis (supporting); Writing – review and editing (supporting). **Su Chin Heo**: Conceptualization (supporting); Writing – review and editing (supporting). **Kwonmoo Lee**: Conceptualization (supporting); Writing – review and editing (supporting). **Andrew T Lombardo**: Writing – review and editing (supporting). **Yongho Bae:** Conceptualization (lead); Investigation (supporting); Supervision (lead); Writing – original draft (lead); Writing – review and editing (lead).

## DATA AVAILABILITY

The data that support the findings of this study are available from the corresponding author upon reasonable request.

